# The role of two-component system response regulator *BceR* in antimicrobial resistance, virulence, biofilm formation, and stress response of group B Streptococcus

**DOI:** 10.1101/340679

**Authors:** Ying Yang, Mingjing Luo, Haokui, Carmen Li, Alison W. S. Luk, GP Zhao, Kitty Fung, Margaret Ip

## Abstract

The hypervirulent Group B Streptococcus (*Streptococcus agalactiae*, GBS) serogroup III clonal cluster 17 has been associated with neonatal GBS invasive disease and meningits. Serogroup III, ST283 has recently been implicated in invasive disease among non-pregnant adults in Asia. These strains cluster with strains from freshwater fishes from aquaculture and a foodborne outbreak of sepsis, especially with septic arthritis, had been linked to such consumption in Singapore in 2015. Through comparative genome analyses of invasive and non-invasive strains of ST283, we identified a truncated response regulator gene in the non-invasive strain. This two component response gene, previously named a DNA binding regulator, is conserved among GBS strains and is a homologue of *Bacillus subtilis BceR*, the response regulator of the BceRSAB system. Loss of function of the *BceR* response gene in the invasive GBS strain demonstrated bacitracin susceptibility in *ΔBceR* mutant with MICs of 256-fold and four-fold reduction in bacitracin and human cathelicin LL-37 compared to wild type and complementation strains. Upregulation of *dltA* of wild type strain vs *ΔBceR* mutant was demonstrated (*p*<0.0001), and was previously shown in *Staphylococcus aureus* to resist and repel cationic peptides through excess positive charges with D-alanylation of teichoic acids on the cell wall. In addition, *ΔBceR* mutant was less susceptible under oxidative stress under H_2_O_2_ stress when compared to wild type strain (*p*<0.001) and inhibited biofilm formation (*p*<0.05 and *p* < 0.0001 for crystal violet staining and cfu counts). The *ΔBceR* mutant also showed reduced mortality as compared to wild type strain (*p*<0.01) in a murine infection model. Taken together, *BceRS* is involved in bacitracin and antimicrobial peptide resistance, survival under oxidative stress, biofilm formation and play an important role in the virulence of GBS.

**Author Summary:** Two-component systems (TCSs) play an important role in virulence in bacteria, and are involved in detecting environmental changes. Although *S. agalactiae* was reported to contain more predicted TCSs than *Streptococcus pneumoniae,* few have been studied in detail. In this work, comparative genomic analysis of GBS invasive (hyper-virulent) and non-invasive serotype III-4 strains were performed to determine any gene differences that may account for severity of disease in humans. *BceR-like* TCS was selected and suspected to be involved in virulence, and thus *BceR* was deleted in a hyper-virulent GBS serotype III-4 strain. We demonstrated that this *BceR-like* TCS is involved in GBS virulence and induced proinflammatory host immune responses. Our study of TCS *BceR* may guide further research into the role of other TCSs in GBS pathogenicity, and further explore therapeutic targets for GBS disease.

## Introduction

Group B streptococcus (GBS) is a leading cause of sepsis in neonates and pregnant mothers worldwide [1, 2]. GBS serogroup III ST 17 has been well associated with hyper-virulence in causing neonatal sepsis and meningitis [3]. In addition, life-threatening syndromes of toxic shock syndrome and meningitis due to GBS are increasingly reported in non-pregnant adults [4]. As in other regions, serotypes I, III, and V are predominant in adult GBS invasive diseases in Hong Kong (HK) [5].

Serotype III-4/ST283 strains have been correlated to invasive diseases in non-pregnant adults in HK [6, 7, 8–10]. Moreover, they have been recently associated with an outbreak of GBS invasive disease in adults in Singapore, and suspected to be due to foodborne ingestion of contaminated freshwater fish (sushi) [10]. Compared with other serotypes identified in non-pregnant adults, GBS serotype III-4 has a significantly higher propensity to cause meningitis and septicaemia, and accounted for over 50% of all GBS meningitis in non-pregnant adults due to serotype III during 1993 to 2012 in HK. This organism is also associated with high mortality and increased summer prevalence, and was implicated in the death of a previously healthy adult with toxic shock-like syndrome in HK [6]. In Singapore, the outbreak of this hyper-virulent GBS serotype III-4 strain was estimated to have started in 2015, and the patients with GBS serotype III-4 infection were younger and developed spinal infection and septic arthritis at a high ratio [10]. Over the last 15 years, GBS serotype III-4 strains remain as a single clone of ST283, possess distinct surface protein genes and mobile genetic elements, and exhibit indistinguishable PFGE fingerprints [6], suggesting that GBS III-4 strains may be hyper-virulent and possess special virulent genetic determinants.

At present, −92 complete GBS genomes are available in the GenBank database (https://www.ncbi.nlm.nih.gov/genome/genomes/186). These genomes revealed that GBS possesses many pathogenici islands encoding virulence genes and transcriptional regulators compared to other streptococci species [12]. Novel regulators associated with GBS pathogenesis have also been identified based on genome analyses [13–15].

In the present study, we conducted whole genome sequencing and comparative genomics analysis of GBS serotype III-4/ST283 invasive and non-invasive strains. A two-component system (TCS) BecR-like regulator *BceR* (GenBank: CU_GBS08_01010) was identified to possess a truncated mutation in the non-invasive GBS strain. TCS are key regulators of bacteria to detect and respond to their environmental challenges, and a number of BceR-like systems have been characterized, which constitute part of the antimicrobial peptide detoxification modules in Firmicutes bacteria (Fig 1)[16–17]. The ciaR of TCS ciaRH was reported to impact biofilm formation in *S. sanguinis* SK36 [18]. WalKR system was found to be involved in controlling cell wall metabolism, and negatively controlled by SpdC in *S. aureus* [19]. The TCS regulator LtdR was found to regulate many bacterial processes in *S. agalactiae* persistence and disease progression [20]. All these studies indicate the importance of TCSs in different species of bacteria. Through whole genome analysis in GBS, multiple TCSs have already been reported. However, the role of these TCSs in GBS pathogenesis is not well understood [21–24].

**Figure 1.**
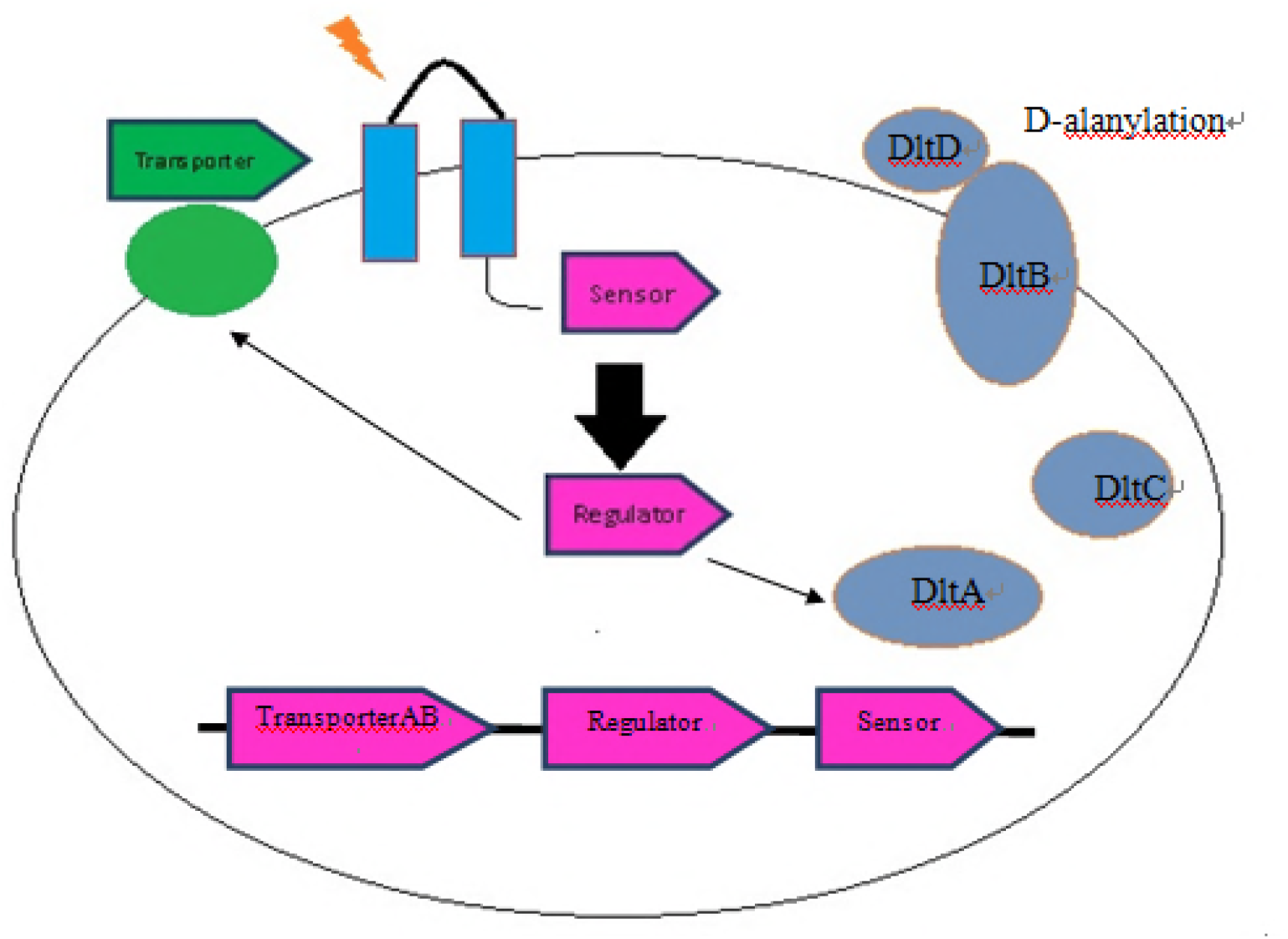
Working model of the antimicrobial peptide sensor and regulator system. The sensor *BceS* is a histidine kinase, which can lead to phosphorylation of BceR. The activated regulator *BceR* can activate the promoter of the transporter *BceAB* to facilitate the resistance. This *BceR* is also found to trigger the expression of dlt operon and result in the D-alanylation of teichoic acids, which in turn decreased negative charge of the bacterial membrane and ensure the resistence.

The best studied example of BceR-like systems is the bacitracin resistance module (BceRS-AB) in *B. subtilis* [17, 25]. *In B. subtilis,* this system is linked with a transporter-*BceAB,* which served as a detoxification pump against peptide antibiotic to monitor antimicrobial peptides (AMPs) and regulate its resistance [17, 25–26]. *S. aureus* was also reported to have two complete TCS/ABC transporter modules, termed *GraRS-VraFG* and *BraRSAB*, whose systems either sense the same type of AMPs or different AMPs, and also interplay with each other to mediate AMPs resistance [17, 27–28].

Apart from AMPs resistance, *GraRS* in *S. aureus* was shown to play an important role in virulence. This TCS also influences autolytic activity and biofilm formation [29]. Transcriptomic study revealed that some major virulence genes or regulators were found to be significantly differentially expressed in GraRS mutant compared with the parental strain [30].

Recently, BceRS-like systems have been deeply characterised into two groups based on detoxification mechanisms: (1) the system that mediates drug-specific resistance, such as MbrAB, in *S. mutans,* by upregulating the expression of the ABC transporter that promotes the removal of antimropeptides; and (2) the system that can not only enhance the expression of ABC transporter, but also lower the overall negative net charge of the cell envelope, such as ApsRS in *S. epidermidis* and GraRS in *S. aureus.* The Aps system is activated by a range of AMPs, and then upregulates the expression of vraF and vraG proteins, which possibly function as an AMP transporter. This Aps system was demonstrated to decrease the anionic charge of the bacterial surface specific to cationic AMPs (CAMPs). In the Aps sensor, negatively charged amino acid residues exist, which can interact with CAMPs. Transduction of the signal occurs through ApsS, and then the AMP resistance mechanisms are activated, which include dlt operon and MprF enzyme, resulting in the decreased negative charge of the bacterial surface [31]. dlt operon was found to encode the proteins necessary for D-alanylation of cell wall teichoic acid, and through the repulsion of the cation, confers resistance to AMPs [32–33]. dltA is a cytoplasmic carrier protein ligase that catalyses the D-alanylation of the D-alanyl carrier protein dltC. dltB is a transmembrane protein, which was reported to efflux activated D-alanine to the site of acylation. dltD is thought to possess multifunctional activities, such as hydrolysis of mischarged dltC [34–35]. We demonstrated here that the *BceR* response regulator is involved in the virulence of GBS III-4 and responds to environmental stress, including cell wall-active antimicrobial peptides and H_2_O_2_ stress. Furthermore, deletion of *BceR* results in significant attenuation of virulence in a GBS murine infection model, and decreased the ability of biofilm formation.

## Results

### Whole genome sequencing and comparative genomics analysis of GBS serotype III-4 strains

The genomes of three invasive and two non-invasive GBS serotype III-4 strains were sequenced by the combined use of Roche 454 and Illimina Solexa Genome Analyser, according to the manufacturer’s instructions, and accepted by GenBank as either draft or complete genomes. The meningitis/septicaemia GBS III-4 genomes were compared against the non-invasive GBS III-4 genomes, and all single nucleotide polymorphisms (SNP) were called using Mauve [36] software from all of the ORFs. Most notably, a regulator was identified in truncation in a non-invasive GBS strain (Table 2). The SNPs were confirmed by PCR-sequencing..

**Table 1.** List of strains in this study.

| GBS Strain | Accession Number | Isolation Site | Clinical Details | Patient Age Group | Sequence Type | Molecular Serotyping Group |
| --- | --- | --- | --- | --- | --- | --- |
| ATCC12403<br>(NEM316) | NC_004368 | Blood | Septicaemia | Infant | 23 | III |
| CU_GBS_00 | JYCT000000000 | Wound | Non-invasive | Non-pregnant adult | 283 | III-4 |
| CU_GBS_08 | CP010874 | Blood | Toxic shock syndrome | Non-pregnant adult | 283 | III-4 |
| CU_GBS_10 | JYCU000000000 | Blood | Septic arthritis | Non-pregnant adult | 283 | III-4 |
| CU_GBS_12 | JYCV000000000 | Vaginal-rectal swab | Non-invasive | Pregnant adult | 283 | III-4 |
| CU_GBS_98 | CP010875 | Cerebrospinal fluid | Meningitis | Non-pregnant adult | 283 | III-4 |
| CU_GBS_08_Δ*BceR* |  |  | *BceR* deletion mutant of CU_GBS_08 |  |  |  |
| CU_GBS_08_Δ*BceR*+*pBceR* |  |  | *BceR* complementation of CU_GBS_08_Δ*BceR* |  |  |  |

**Table 2.** List of truncated genes in non-invasive strain (CU_GBS_12) vs. invasive strain (CU_GBS_08).

| Gene | Mutations in CU_GBS_12 | Potential Function of Protein | References |
| --- | --- | --- | --- |
| *BceR* | c.288delG | - Regulates *bceABRS* operon, required for bacitracin resistance in *Streptococcus mutans* | [16] |
| *mecA* | c.8_20del<br>TGAAACAAATCAG | - Negative regulator of competence in *streptococcus mutans* | [37] |
| *murAB* | c.894delA | - Catalyses first step of peptidoglycan biosynthesis<br>- Resistance to fosfomycin in *Streptococcus pneumoniae*, *mycobacterium tuberculosis*, *chlamydia trachomatis* | [38] |
| *wfgD* | c.110G>A | - Confers resistance to bacitracin in *Streptococcus mutans* | [39] |

### *ΔBceR* mutant strain was more sensitive towards bacitracin and antimicrobial peptides

The MICs of the antimicrobial peptide (AMPs) bacitracin and human cathelicidin LL-37, cell-wall and other classes of antibiotics to the *ΔBceR* mutant, complementation, and wild type strains were tested. The MICs of the mutant strain was 256 and four-fold lower for bacitracin and LL-37, respectively, compared to wild type strain. The *ΔBceR* mutant did not become more sensitive towards colistin and other antibiotics. To demonstrate the specificity of these phenotypes, expression of *BceR* was restored to *ΔBceR* mutant strain using plasmid pDL289. Thus, complementation of *ΔBceR* with pDL289-BceR reversed the antimicrobial sensitivity phenotypes in the presence of bacitracin and LL-37 (Table 3).

**Table 3.** Minimal inhibitory concentrations (MIC) of antimicrobial peptides and other antibiotics in GBS strains.

| Antibiotics | MICs ($\mu\text{g/ml}$) | | | | |
| --- | --- | --- | --- | --- | --- |
|  | CU_GBS_08 | CU_GBS_08_Δ*BceR* | CU_GBS_08_Δ*BceR*+*pBceR* | CU_GBS_12 | ATCC49619 |
| **Bacitracin** | 64 | 0.25 | 64 | 0.25 | 8 |
| **LL-37** | 256 | 64 | 256 | 128 | 2 |
| **Colistin** | 256 | 512 | 256 | 256 | 256 |
| **PolymixinB** | 128 | 128 | 128 | 64 | 128 |
| **Ampicillin** | ≤0.06 | ≤0.06 | ≤0.06 | ≤0.06 | ≤0.06 |
| **Cefotaxime** | ≤0.06 | ≤0.06 | ≤0.06 | ≤0.06(S) | ≤0.06 |
| **Penicillin** | ≤0.06 | ≤0.06 | ≤0.06 | ≤0.06 | ≤0.06 |
| **Vancomycin** | 0.5 | 0.5 | 0.5 | 0.5 | 0.5 |
| **Erythromycin** | ≤0.06 | ≤0.06 | ≤0.06 | ≤0.06 | ≤0.06 |
| **Ciprofloxacin** | 0.5 | 0.5 | 0.5 | 0.5 | 0.5 |

### The gene expression of dltA was reduced in BceR mutant GBS strain

As shown in Fig 2 (A), the gene expression of dltA of GBS strains in THB only were used as a control and adjusted to 1, respectively. The expression of dltA was higher in THB containing Bac at 1/8 MIC for CU_GBS_08 (Bac MIC 64 μg/ml) than that in THB containing Bac at 1/8 MIC for CU_GBS_ΔBceR (Bac MIC 0.25 μg/ml) (p<0.0001), and for CU_GBS_12 (Bac MIC 0.25 μg/ml) (p<0.0001), respectively. However, no significant difference of gene mprF expression was found between wild type strain and BceR mutant strain under treatment with bacitracin. Bacterial survival rate under of H_2_O_2_ stress was decreased in ΔBceR mutant strain.

The *ΔBceR*, wild type, and complementation strains in response to H_2_O_2_ stress were measured (Fig 3). The mutant strain was significantly more sensitive to be killed by H_2_O_2_ than wild type strain. The survival rate of mutant strain was approximately 20% less than wild type strain (p<0.001), although no statistically significant difference was observed between wild type and non-invasive CU_GBS_12 strain.

**Figure 2.**
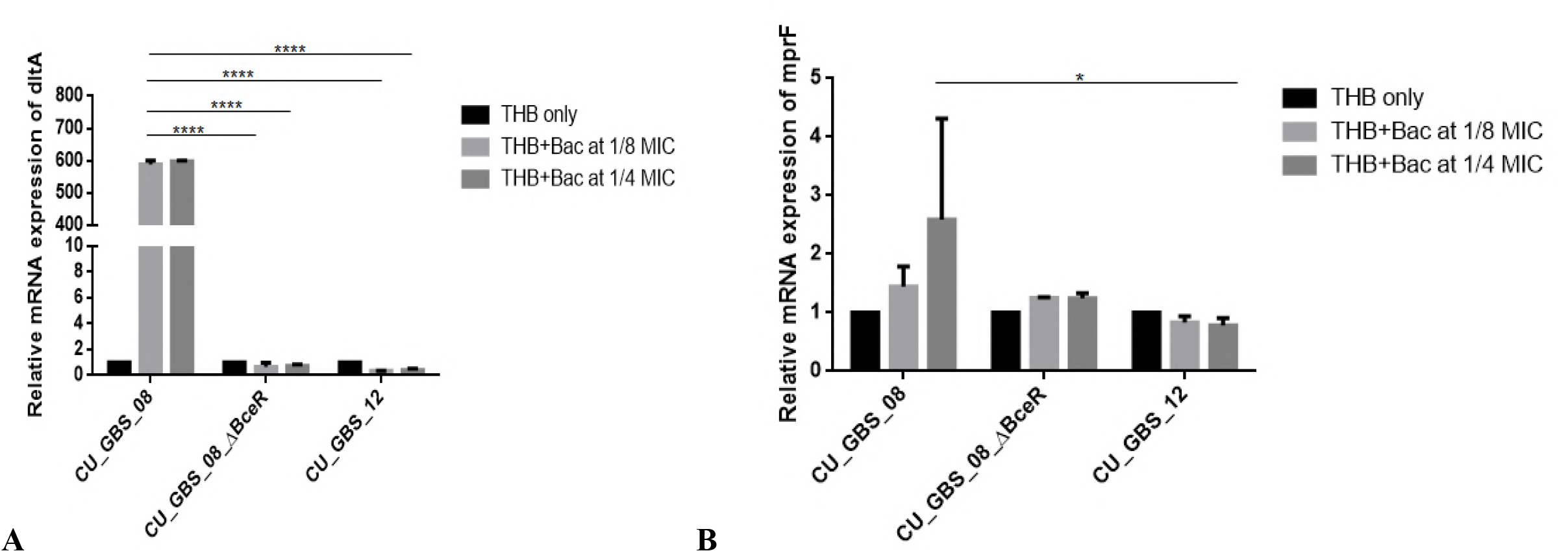
Relative expression of dltA and *mprF* in *S. agalactiae* strains with addition of bacitracin (Bac) (A-B). Gene expression of *dltA* and *mprF* in wild type GBS III-4 strain, CU_GBS_08 (Bac MIC 64 μg/ml), two-component system deletion mutant (CU_GBS_08_ΔBceR, Bac MIC 0.25 μg/ml), and non-invasive GBS III-4 strain CU_GBS_12 (Bac MIC 0.25 μg/ml) were shown. GBS strains without treatment were normalized to 1. Error bars represent the standard deviation of the mean values from at least three replicates. Significance was determined by one-way ANOVA (\**p* < 0.05 and \*\*\*\**p* < 0.0001).

**Figure 3.**
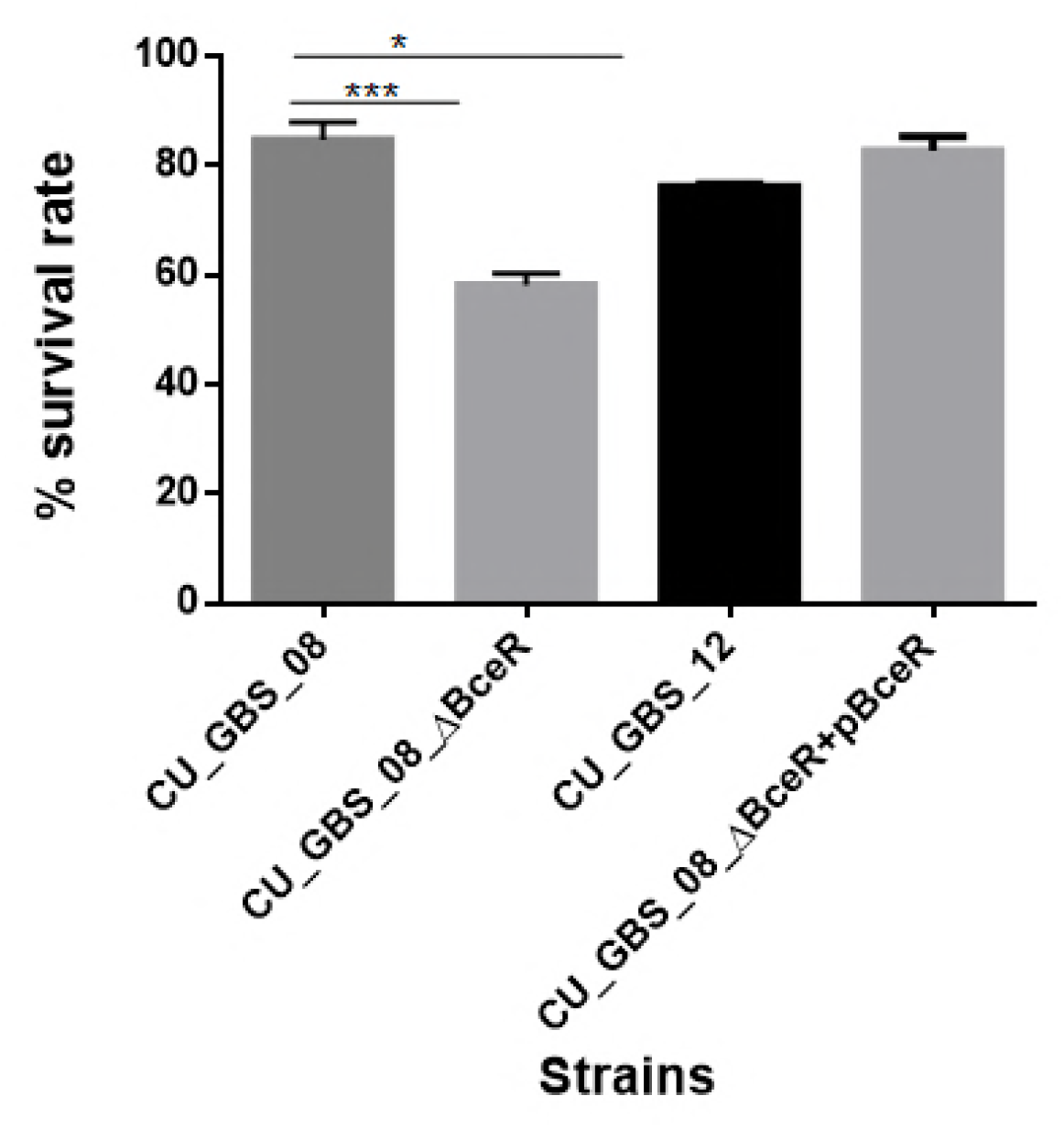
Effect of H_2_O_2_ stress on GBS. The *ΔBceR* was significantly more susceptibility to H_2_O_2_ (40 mM) exposure than wild type (\**p* < 0.05 and \*\*\**p* < 0.001).

### *ΔBceR* mutant strain showed impaired ability in biofilm formation

Four GBS strains, wild type, *ΔBceR* mutant strain, *ΔBceR* complementation, and CU_GBS_12 (non-invasive strain) were examined to assess the effect of *BceR* on biofilm formation by crystal violet staining and cfu counting, respectively (Figs 4 and 5). One-way ANOVA analysis showed that biofilm formation was impaired significantly in *ΔBceR* mutant when compared with wild type strain (*p*<0.05 and *p*<0.0001 for crystal violet staining and cfu counting, respectively) and was reversed by the complementation strain.

**Figure 4.**
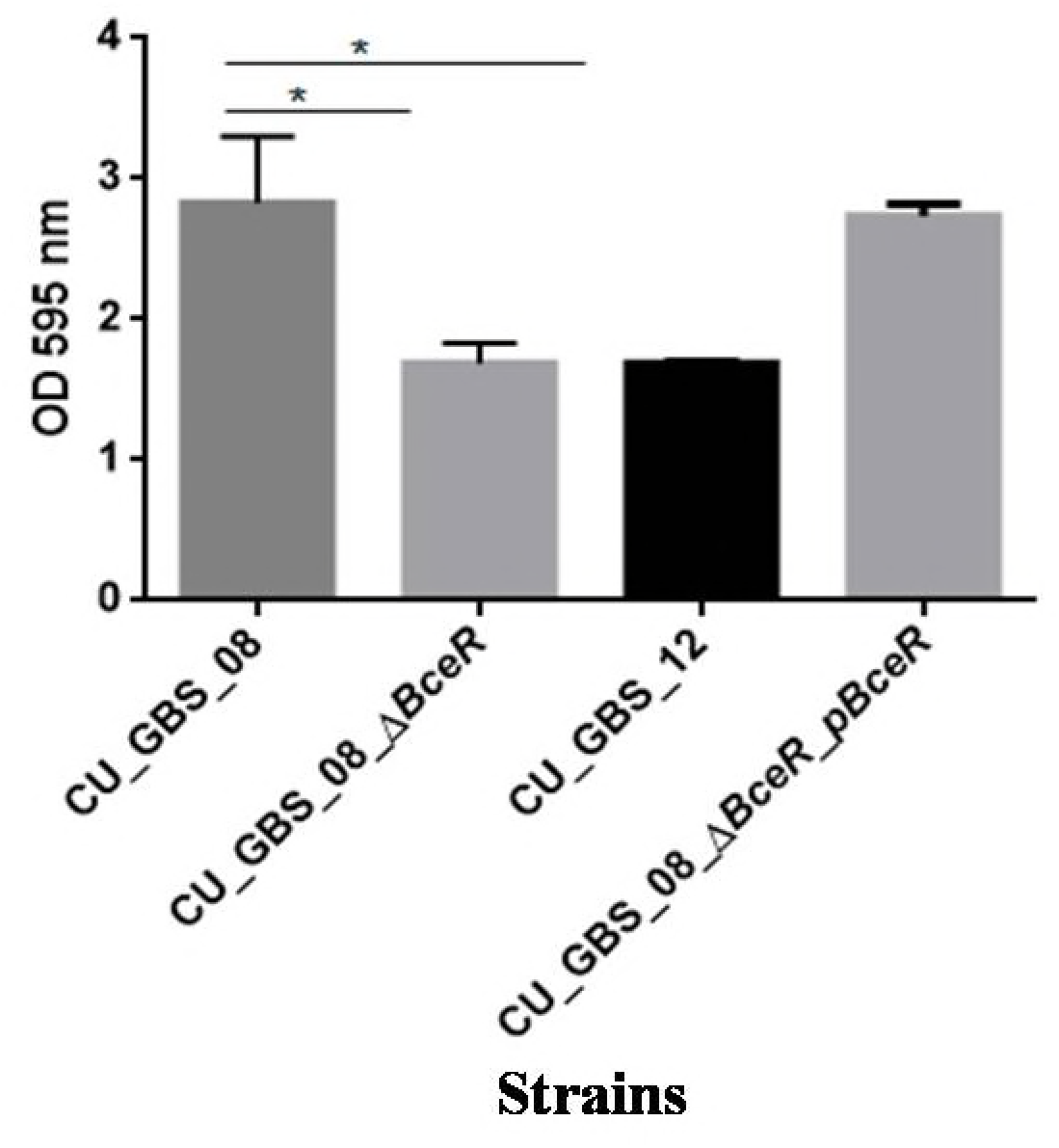
Effect of two-component regulator *BceR* on biofilm formation in *S. agalactiae* serotype III-4. Bacteria stained with crystal violet and measured at OD_595_. Significance was determined by one-way ANOVA (\**p* < 0.05).

**Figure 5.**
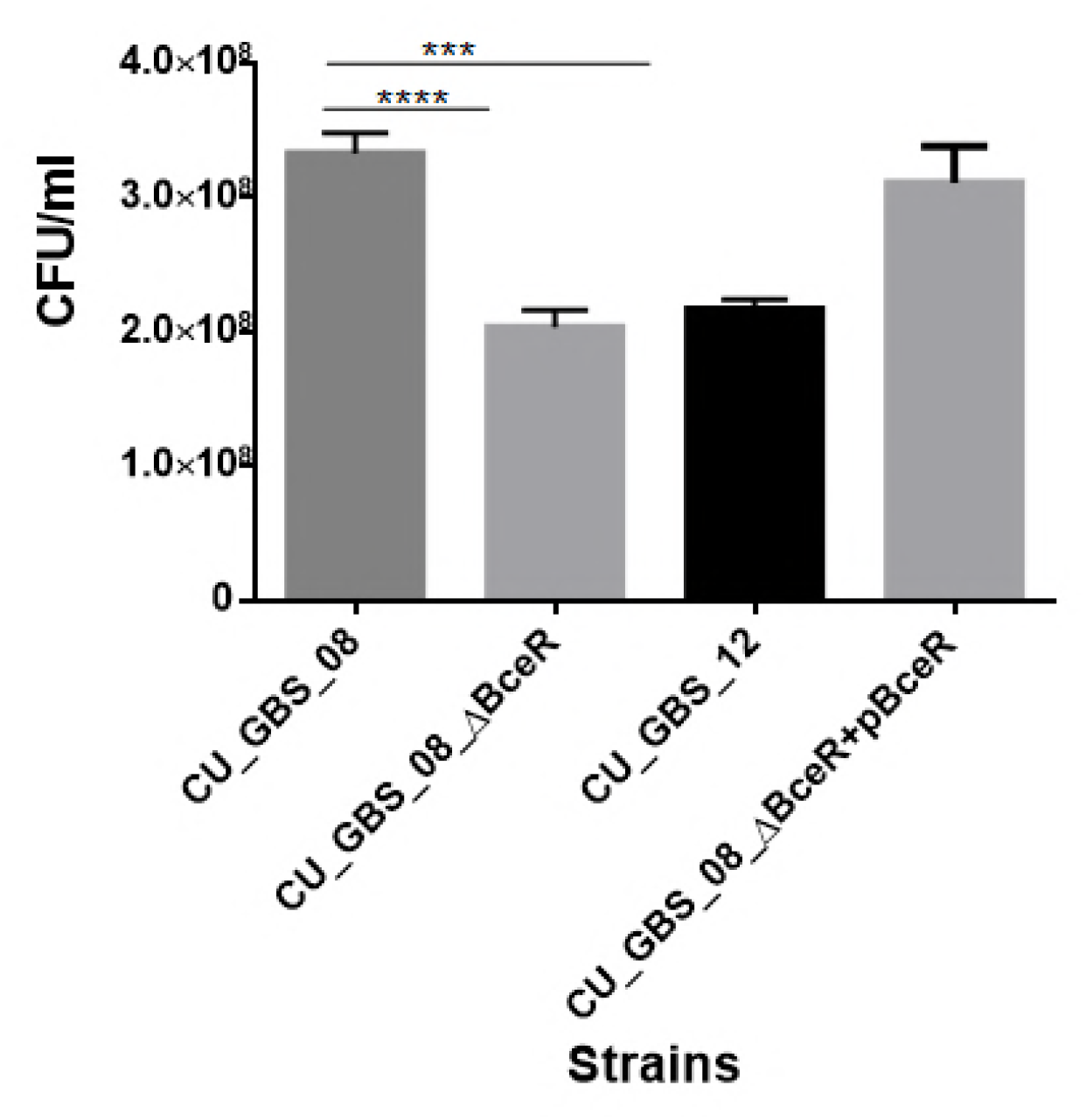
Bacterial counts (CFU) of biofilms grown in *S. agalactiae* serotype III-4 wild type and mutant strains. Significance was determined by one-way ANOVA (\*\*\**p* < 0.001 and \*\*\*\**p* < 0.0001).

The biofilms were also evaluated with confocal microscopy (CLSM), and cell density (xy images) and thickness (xz images) of biofilms were assessed. In the images taken by the microscopy, as shown Fig 6 (A-C), most cells in the biofilms were stained in green, indicating that more live cells were present. Less signal was captured from the xy image and xz image of *BceR* mutant strain when compared with wild type strain, which indicated that few living or dead cells presented in the *ΔBceR* mutant strain. Images taken with CLSM revealed that loss of BceR-like regulator possibly inhibited the formation of biofilm, resulting in lower cell density and reduced thickness.

**Figure 6.**
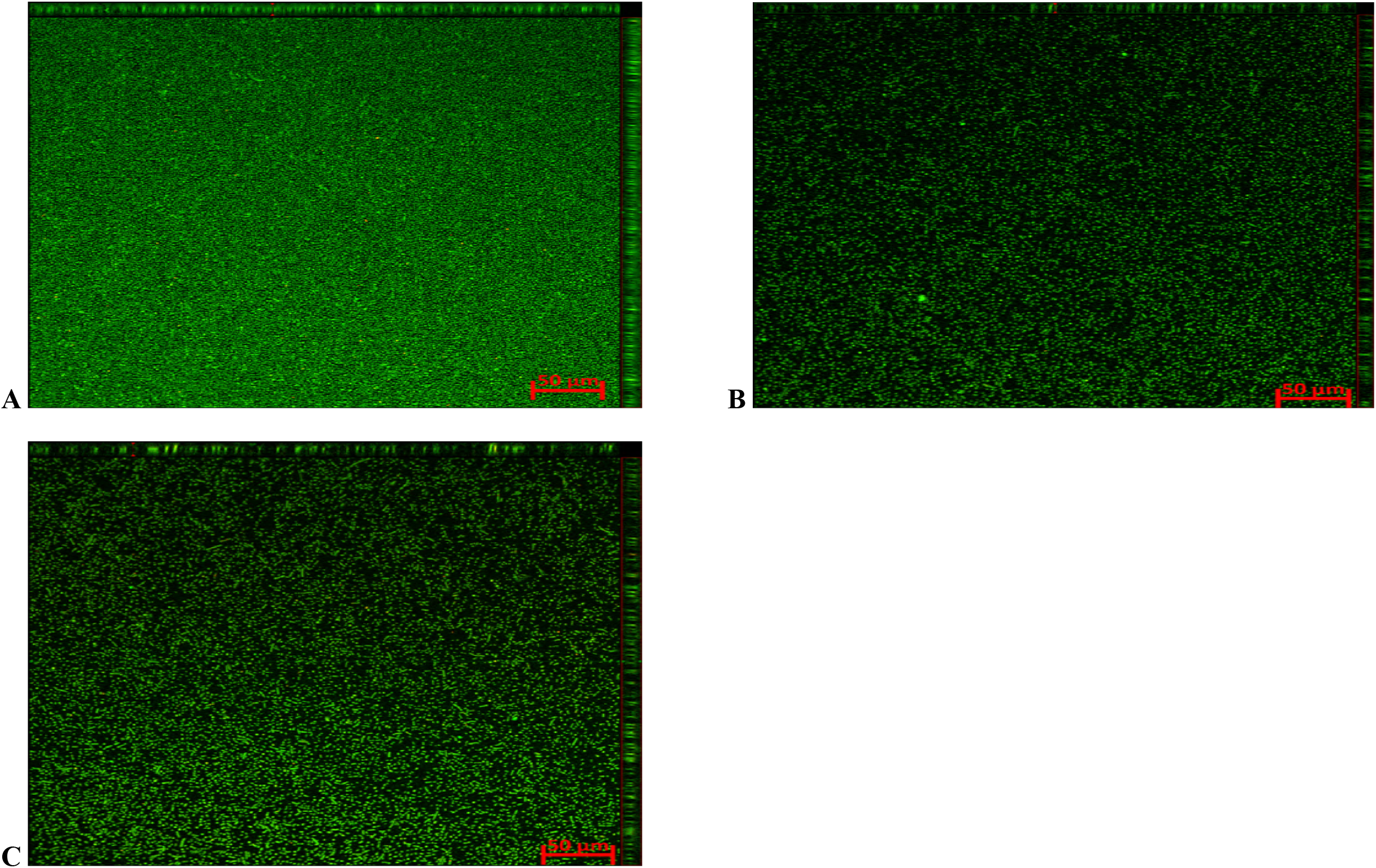
Confocal microscopic images of biofilms in CU_GBS_08, CU_GBS_08_*ΔBceR*, and CU_GBS_12 strains, respectively (A-C). Biofilm formation was inhibited in *ΔBceR* strain. Biofilms were stained with BacLight, showing viable (green fluorescence) and nonviable (red fluorescence) GBS bacteria within the biofilms. The assays were performed at least three times.

### Mitogenicity and pro-inflammatory response induced by GBS in human PBMCs

Proliferation of PBMCs was evaluated after 24 h of stimulation with GBS strains or PHA (10 μg /ml) by measuring [3H] thymidine incorporation. As shown in Fig 7, although all bacteria induced proliferation of PBMCs, *ΔBceR* demonstrated a statistically significant difference in the reduction ability of immunogenicity (*p*<0.0001). Similarly, the cytokine levels of TNF-α, IL-6, IL-8, IL-1β, IL-10, and IL-12 were determined, as shown in Fig 8A-F. Statistically decreased expression of pro-inflammatory cytokines were obtained when compared between the wild type strain and the isogenic mutant strain *ΔBceR*. The decreased release of TNF-α was more obvious (*p*<0.0001), approximately four-fold lower than the wild type strain. This was followed by IL-6, IL-1β and IL-10, approximately a two-fold decrease in *ΔBceR* mutant strain (*p*<0.001 for IL-6, IL-10, and *p*<0.01 for IL-1β). The peak of IL-6 expression was delayed to 24 h in the *ΔBceR* mutant strain, and the release of IL-8 was approximately 1.4-fold lower for the *ΔBceR* mutant strain (*p*<0.0001). The release of IL-12 could not be detected in both wild type and mutant strain. Complementation of *ΔBceR* with pDL289 reversed the cytokines release.

**Figure 7.**
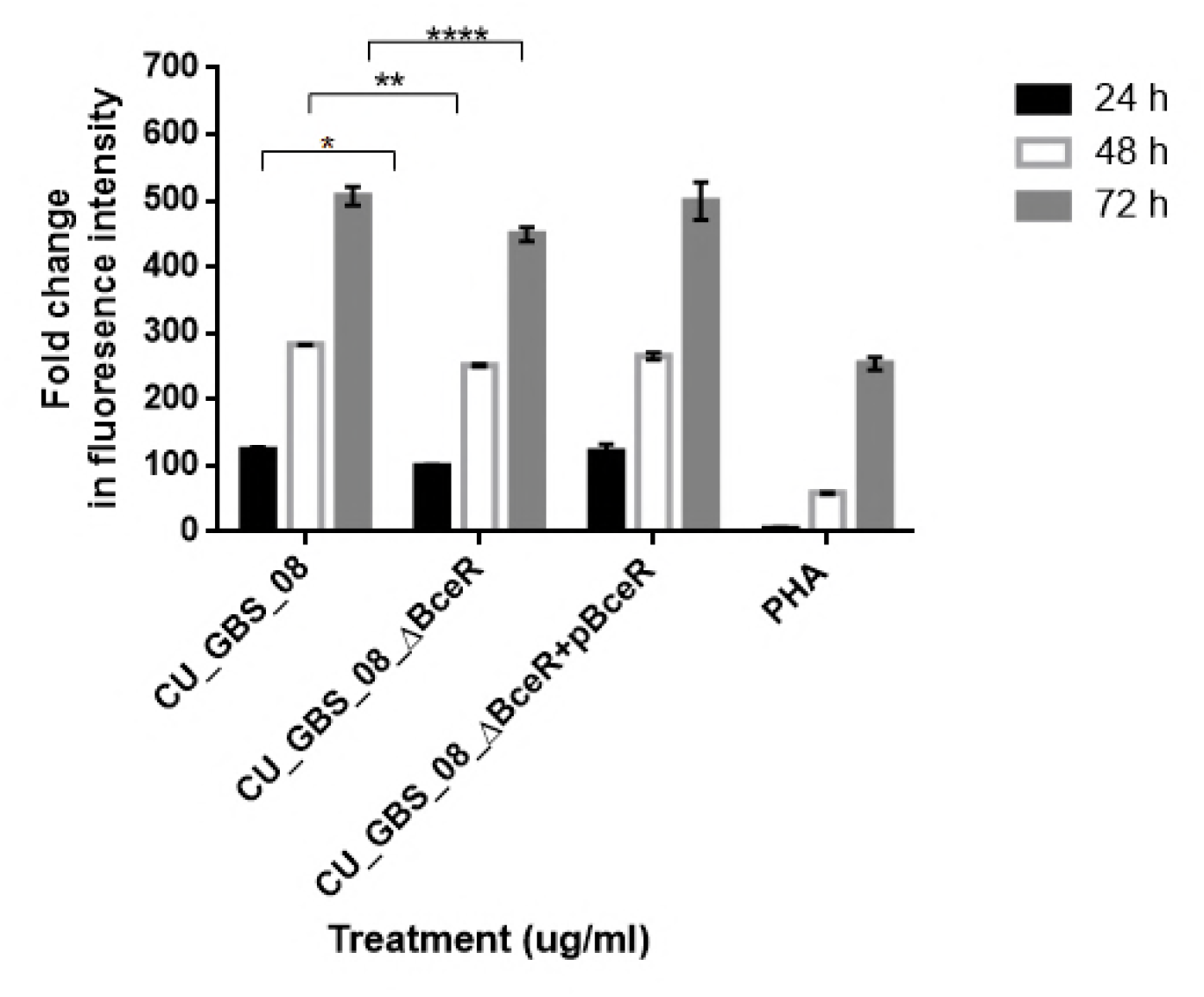
Effect of *BceR* deletion on proliferation of PBMCs. PBMCs were cultured in 25 μg/ml heat-killed GBS strains for 24 h, with phytohaemaglutinin (PHA) (10 μg/ml) only or medium only as controls. Cell proliferation was determined by fluorescence intensity. Three independent experiments were performed. Data are expressed as mean ± SD. Statistical significance at * p < 0.05, ** p < 0.01, and **** p < 0.0001 were reached when GBS wild type strain was compared to the *BceR* deletion mutant (CU_GBS_08_ΔBceR).

**Figure 8.**
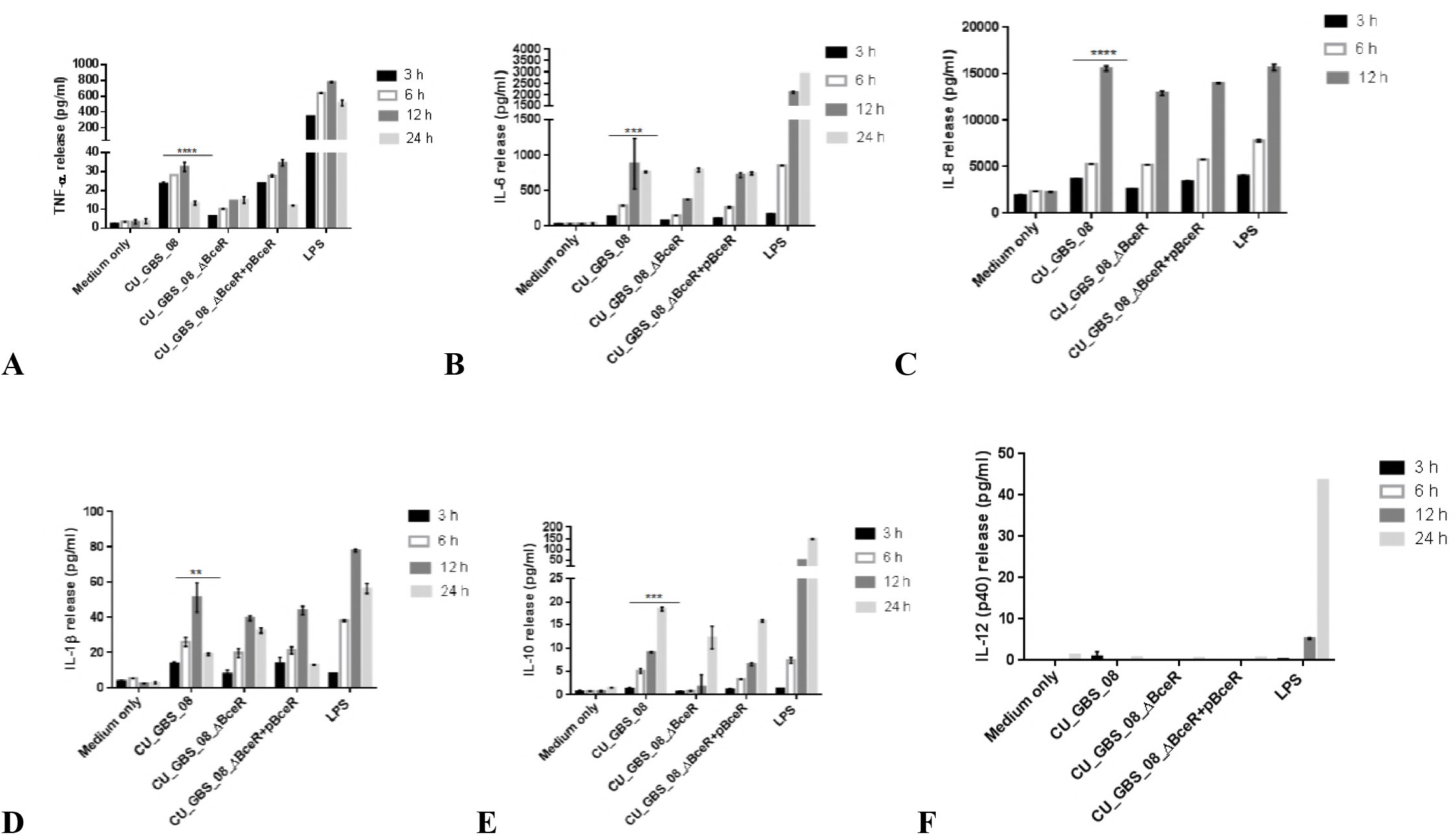
Release of TNF-α, IL-6, IL-8, IL-1β, IL-10, and IL-12 from human lymphocytes as induced by wild type GBS and *ΔBceR* mutant. Heat-killed GBS extract (5 μg/ml) (invasive strain CU_GBS_08; CU_GBS_08_*ΔBceR*, two-component system deletion mutant; CU_GBS_08_*ΔBceR+pBceR*, *BceR* complemented strain) was incubated with human lymphocytes. Cytokine release was measured at 3 h, 6 h, 12 h, and 24 h. LPS represents the lipopolysaccharide control. Data are expressed as mean ± SD (** p < 0.01, *** p < 0.001, and **** p < 0.0001).

### Deletion of *BceR* resulted in attenuation in virulence in a mouse infection model

The virulence of this GBS III-4 wild type strain was compared to *ΔBceR* mutant strain in an intraperitoneal-injection mouse model. *ΔBceR* mutant strain had a LD_50_ of 1*10^7^CFU, which is nearly one order of magnitude higher than wild type strain (3*10^6^ CFU) (Table 4). The survival rate of mice infected with GBS at10^^7^ CFU I.P. after 10 d of inoculation is shown in Fig 9. The *ΔBceR* mutant was attenuated compared to the wild type strain, and revealed an increased survival rate of 23.3% versus 0% in the wild type strain.

**Figure 9.**
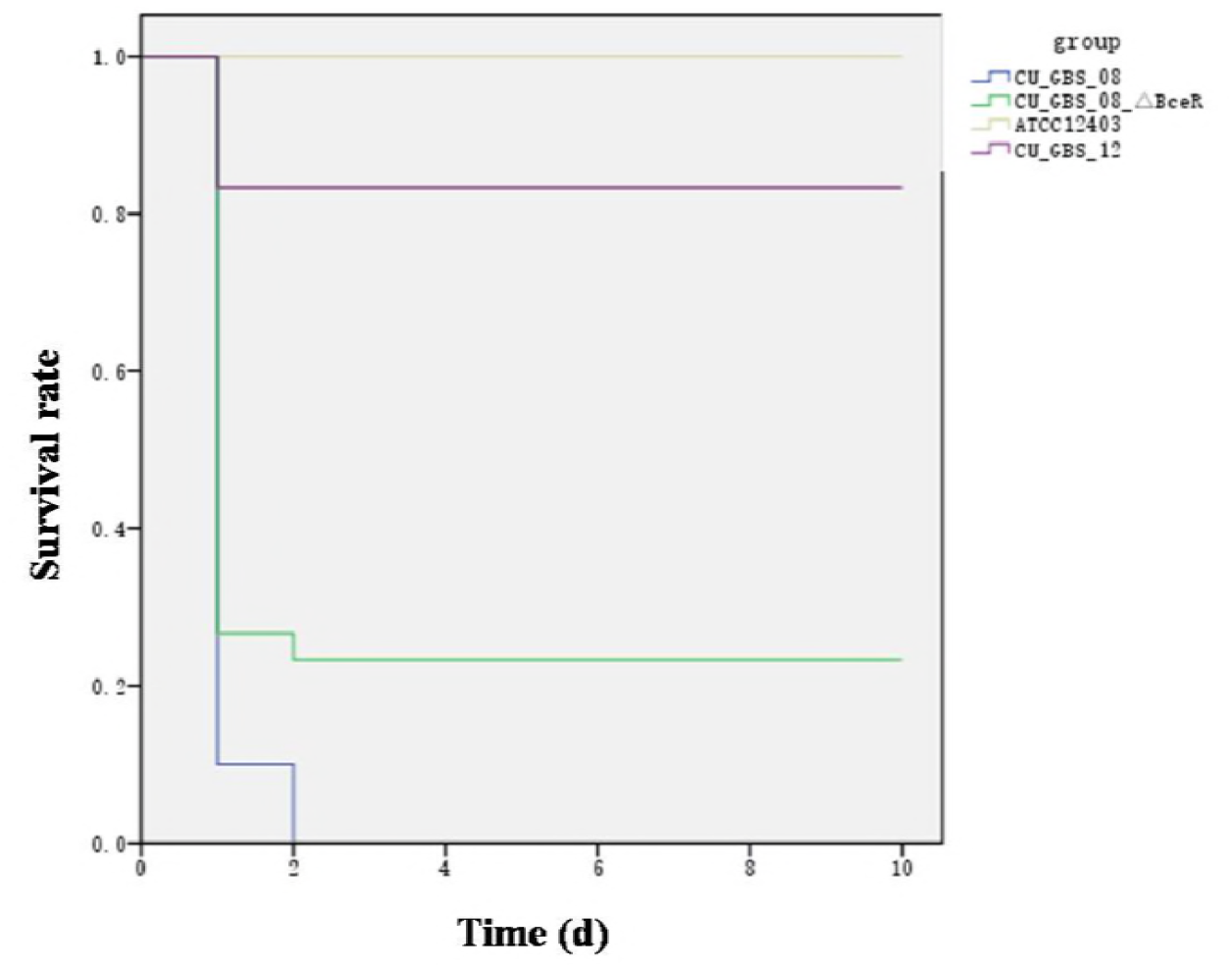
Kaplan-Meier survival curve.

**Table 4.** LD_50_ of GBS infection murine models.

| GBS Strain | LD<sub>50</sub><sup>a</sup> |
| --- | --- |
| CU_GBS_08 | $3 \times 10^6$ |
| CU_GBS_08_Δ*BceR* | $1 \times 10^7$ |
| CU_GBS_12 | $2 \times 10^7$ |
| ATCC 12403 | $3 \times 10^8$ |

<sup>a</sup>LD<sub>50</sub> expressed as bacterial c.f.u. (colony-forming units). LD<sub>50</sub> calculated by survival rate 10 d post-intraperitoneal injection, using the method of Reed and Muench (1938).

### Deletion of *BceR* resulted in difference of protein expression of GBS strain

Proteomic analysis of *S. agalactiae* using two-dimensional electrophoresis and mass spectrometry showed that 3 proteins have significantly decreased expression in *ΔBceR* mutant strain (>2 folds reduction in expression), and boolean operation was used for comparation of the intensity of the expression of protein spots (Table 5, Fig 10). The protein Alkyl hydroperoxide reductase (AhpC) revealed a 2.72 folds change, and this protein was reported to be induced by oxidative stress and positively controlled by the *oxyR* gene in *Salmonella typhimurium.* The Gls24 family stress protein (Gls24), and Alcohol dehydrogenases (ADH) had 2.79 and 2.59 folds change respectively. Realtime PCR was conducted to confirm the results of 2DE-mass spectrometry at RNA level, and the three proteins revealed 6.73, 3.56 and 6.7 folds reduction at RNA level in *ΔBceR* mutant strain respectively (Fig 11).

**Figure 10.**
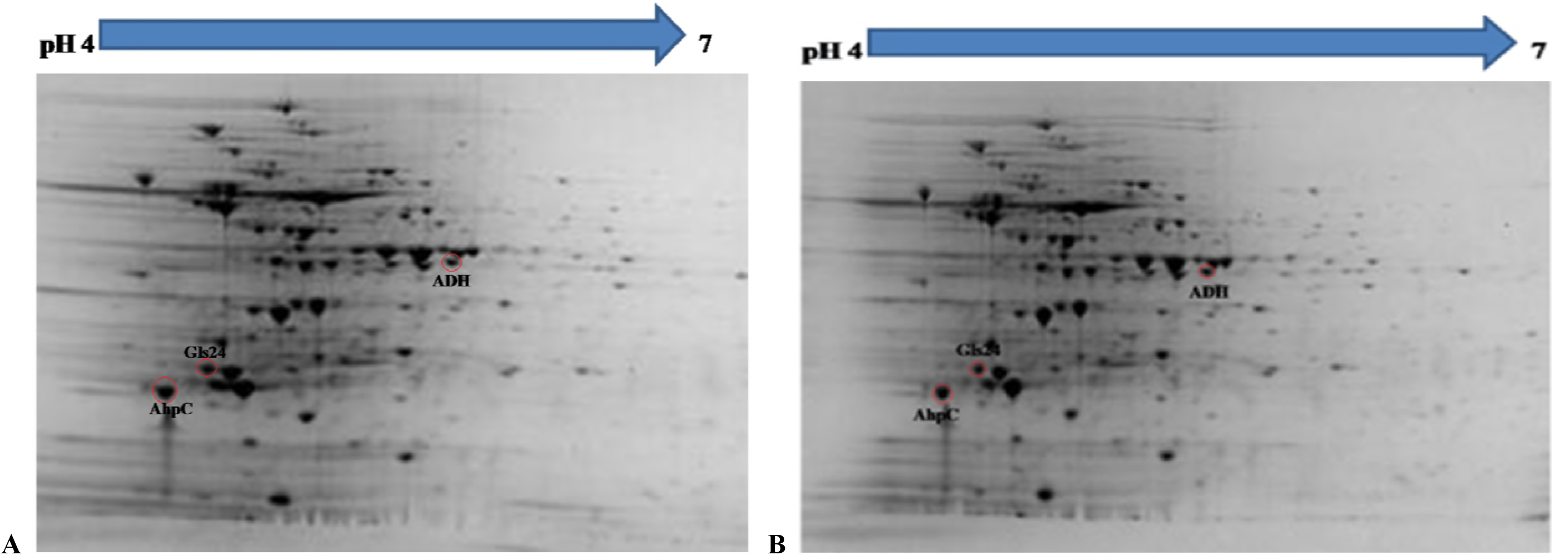
Photographs of 2-DE gel in wild type and *ΔBceR* mutant GBS strain. Red cycles showed the proteins that found to have significantly decreased expression in *ΔBceR* mutant strain (A-B). They were identified as Alkyl hydroperoxide reductase (AhpC), The Gls24 family stress protein (Gls24), and Alcohol dehydrogenases (ADH) respectively by mass spectrometry. The gel photos were normalized and compared using software PDQuest (Version8.0.1, Bio-Rad, USA), Boolean method was chosen for detecting the proteins with statistic significantly difference in expression between GBS III-4 wild type and mutant strains.

**Figure 11.**
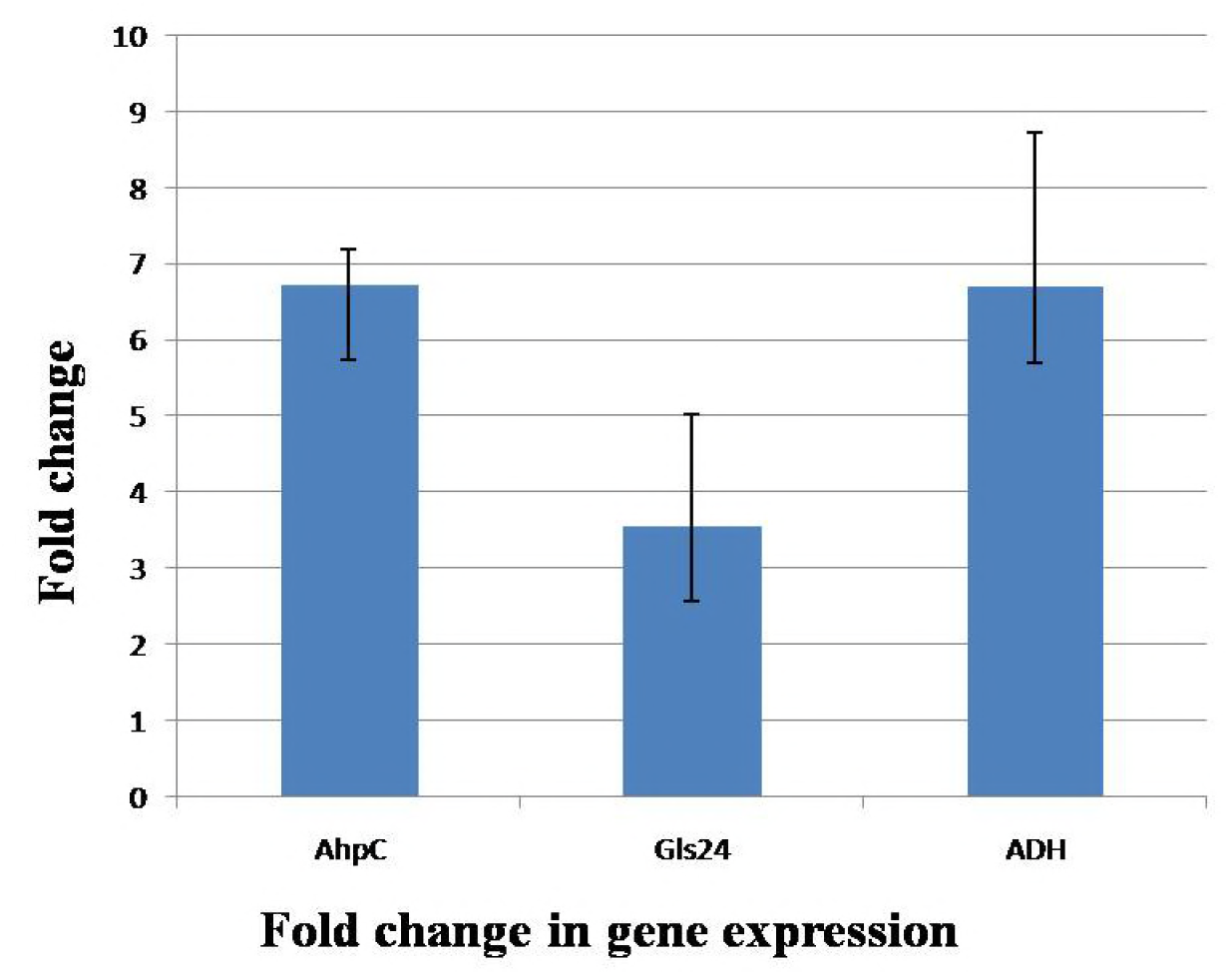
Application of the 2^-ΔΔCT^ method. The experiment was conducted to validate the effect of *BceR* gene knockout on the expression of candidate genes. *AhpC,* alkyl hydroperoxide reductase; *Gls24,* Gls24 family general stress protein; *ADH,* zinc-dependent alcohol dehydrogenase. Error bars represent the standard deviation of the mean values from at least three replicates.

**Table 5.** Results of proteins identification by mass spectrometry.

| Spot | Protein name | Protein score | Database | Accession No | References |
| --- | --- | --- | --- | --- | --- |
| 1 | Alkyl hydroperoxide reductase (AhpC) | 391 | NCBIInr | gi\|403643060 | [40] |
| 2 | Gls24 family stress protein (Gls24) | 291 | NCBIInr | gi\|446353446 | [41] |
| 3 | Alcohol dehydrogenases (ADHs) | 409 | NCBIInr | gi\|446571849 | [42-43] |

## Discussion

Through whole genome sequencing and comparative genome analysis of invasive and non-invasive group B Streptococcus (GBS) Group III-ST283 strains, a non-synonymous substitution of a DNA binding regulator leading to a truncation of the gene was identified. BLAST analyses revealed that this regulator exist in all GBS strains and was most closely related to the two-component response regulator BceR in *Streptococcus gallolyticus,* with 69% protein sequence homology (GenBank no: CDO17747.1), so we named this regulator as *BceR,* however, this regulator still harbours around 30% difference compared with *BceR* in other bacteria, which means that this two component response regulator possibly get specially founction(s) in GBS. This study revealed that loss of BceR-like regulator in GBS resulted in increased sensitivity of *S. agalactiae* to bacitracin and human cathelicin LL-37. This concurred with a previous study that showed the BceR-like system is related with resistance to cell wall antimicrobial peptides in *Bacillus subtilis* [26, 44]. The GraRS system in *S. aureus* was previously found to respond to vancomycin and polymixin B, and the homologous proteins were determined to mediate resistance to bacitracin and nisin in *S. mutans* and *L. lactis,* respectively [45–46]. The regulation of transporter AB, which can pump out the AMPs from the bacterial cells, is the major mechanism of AMPs resistance of the BceR-like system. However, the loss of GBS *BceR* could not change GBS sensitivity to erythromycin and beta-lactam antibiotics [27–28]. These results demonstrate that the structurally homologous BceRS system may play a unique role in GBS, and highlights the importance of determining the individual effect of the BceR-like system in the pathogenesis of disease in different Gram-positive species.

It was found that in S. *epidermidis,* except genes coding for the ABC transporter, the LTA alanylation system dltAB and the lipid modification enzyme MprF were also reported to be under the control of the BceR-like system [47–48]. In *Listeria monocytogenes,* VirRS was also determined to regulate an array of processes, including adhesion and entry into eukaryotic cells. In addition, the set of genes under control of VirRS includes dltABCD operon, *mprF* gene, and ABC transporter. In *S. agalactiae,* D-alanylation of lipoteichoic acids was found to confer resistance to cationic peptides, and the deficient of dltAB was related to increased sensitivity towards phagocytic cells and attenuated bacterial virulence [49–50]. We found that the expression of *dltA* was upregulated in wild type GBS strain at 1/8 MIC of bacitracin treated. However, there was no statistically significant difference in the expression of *mprF* gene between wild type and *ΔBceR* mutant strain, thus this *dltA* could possibly be a reason for part of the functions of *BceR* in GBS. The formation of D-alanyl-lipoteichoic acid is also found to be associated with adhesion and virulence of *Listeria monocytogenes* [51–52].

From the results of 2D-mass spectrum, the proteins alkyl hydroperoxide reductase-AhpC, zinc-dependent alcohol dehydrogenase-ADH, and gls24 family protein werefound decreased expression in the *ΔBceR* mutant strain, and thehe proteins were reported to be related with with oxidative stress and biofilm formation [40]. Thus, this BceR-like regulator may be involved in the resistance of oxidative stress in *S. agalactiae* serotype III-4. Based on these and the specially living condition of GBS, the environmental stress response ability of wild type/ mutant pairs were evaluated and the *BceR* mutant strain displayed increased sensitivity to H_2_O_2_ stress when compared with wild type strain, which is also in line with the report in TCS GraRS of *S. aureus,* which is involved in the resistance to superoxide radicals [53]. The survival rate of wild/mutant pairsunder different pHs, temperatures, and osmotic pressures were also detected. It survived poorly under low pH (<5), which is consistent with results from other studies of GBS [54]. However, statistically significant differences in pH tolerance, temperature tolerance, and osmotic stress resistance between wild type and *ΔBceR* mutant strains were not detected (data not published).

Bacterial cells within biofilms are difficult to eradicate, as they are highly resistant to antibiotics and the host immune system. The difference in biofilm-forming ability between GBS isolates from asymptomatic pregnant women carriers against clinical infections was found to be statistically significant [55]. The protein Alcohol dehydrogenases (ADHs) was reported to catalyze the reversible conversion of acetaldehyde to ethanol, the ethanol is known to enhance the production of *Staphylococcus* biofilm, and ADH1 expression is known to be upregulated in *Staphylococcus* biofilms [42–43]. In our study, all of the strains were able to form biofilms, and the biofilm biomass of wild type strain is significantly higher than the *ΔBceR* mutant strain. This is consistent with the finding of TCS GraRS in *S. aureus*, which was related to bacterial biofilm formation capacity [29].

The ability of *S. agalactiae* serotype III-4 wild type and *ΔBceR* mutant strain in activating lymphocytes and cytokines production was evaluated in this study. *S. agalactiae* was reported to escape from the host immunity and express immunomodulatory proteins [56]. In our study, we found that the *ΔBceR* mutant strain reduced the levels of lymphoproliferation. It was also determined that the expression of IL-8, a major activator of neutrophils and lymphocytes [57–59], was reduced. It was reported that the BceR-like system mutant strains were more sensitive to be killed by neutrophils in *S. epermidis* and *S. aures* [60]. Cytokines are soluble proteins that play a significant role in inflammatory and immunoregulation of immune responses [57]. In this study, cytokine TNF-α was detected at 3 h after infection of GBS wild type strain, and had a higher expression ratio compared with *ΔBceR* mutant strain. The same condition was found in the expression of IL-6 and IL-1β, and these three cytokines were reported to be positively related with disease severity [57, 59, 61]. Expression of IL-12 could not be detected in this study, and accordingly there was an increased release of IL-10, which was reported to have a suppressive effect on immunity. This is the opposite against IL-12, and may constitute an adaptive response of host immunity towards increased lymphoproliferation and related cytokine expression [62]. Our results demonstrated the mitogenicity nature of this regulator and its ability to induce significant pro-inflammatory cytokines that characterize the development of sepsis and septic shock. The deletion of *BceR* gene did not completely abrogate the proliferation of mononuclear cells and the cytokines release, suggesting other factors are also involved in the virulence and pathogenicity of this strain. An important finding of this study is the link between the BceR-like system and GBS virulence. An attenuation of the *ΔBceR* mutant strain was demonstrated. The virulence of this GBS III-4 wild type strain was compared to *ΔBceR* mutant strain, and we found that the *ΔBceR* mutant strain had a LD_50_ of 1 *10^7^CFU, it is nearly one order of magnitude higher than wild type strain (3*10^6^ CFU) (Table 4). The mortality rate at 10^^7^ CFU i.p. after 10 days was also calculated to further evaluate the virulence of the *ΔBceR* mutant strain, and the *ΔBceR* mutant was attenuated compared with the wild type strain, and revealed increased survival of 23.3% versus 0% in the wild type strain. This is consistent with the role of the BceR-like system in organism virulence, in which TCS is involved in the regulation of numerous virulence factors in *S. aureus* and *L. monocytogenes* [63–64]. Further studies are needed to elucidate this issue.

In this study, potential virulent genes were listed in GBS III-4 for further study. Our results demonstrated the effects of *BceR* on virulence, biofilm formation, and H_2_O_2_ stress response in the GBS strain. It was found that TCS may constitute an effective target for GBS combination therapy [65].

## Material and methods

### Ethics Statement

Animal experiments were conducted at The Laboratory Animal Services Centre in compliance with International Guiding Principles for Biomedical Research Involving Animals and The Hong Kong Code of Practice for Care and Use of Animals for Experimental Purposes. Protocols were approved by the University Animal Experimentation Ethics Committee (AEEC) (Reference no.: 13-063-MIS).

The study was approved by the clinical research ethics committee of the Joint University of Hong Kong-New Territories East Cluster (CRE-2012.054). GBS strains were isolated from patients admitted to a university hospital in Hong Kong, and all samples were anonymized.

### Bacterial strains and growth conditions

Five *S. agalactiae* GBS III-4 clinical strains were originally obtained from Prince of Wales Hospital (PWH). GBS strains were grown in Todd–Hewitt broth (THB) or THY broth (THB supplemented with 5 g/L yeast extract) or on THY blood agar plates (all from Difco Laboratories). Recombinant DNA manipulations were performed in E. coli strain XL-Blue, grown at 37°C in Luria–Bertani (LB) broth (Difco Laboratories) or on LB agar plates.

### Whole genome sequencing and comparative genomics of five GBS serotype III-4 strains

Five GBS strains of serotype III subtype 4 and sequence type ST283 were selected for genome sequencing (CU_GBS_00, CU_GBS_10, CU_GBS_12, CU_GBS_98, and CU_GBS_08). These strains were isolated in Hong Kong (HK) between 1998 and 2012, from both invasive and non-invasive sites in adult patients. Genomic DNA from the GBS strains was extracted with the Wizard^®^ Genomic DNA Purification Kit according to the manufacturer’s protocol for Gram-positive bacteria (Qiagen, Limburg, Netherlands). Genomes were assembled using the metAMOS pipeline (version 1.5rc3) [66]. The draft genomes of CU_GBS_00, CU_GBS_10, and CU_GBS_12 were deposited in the NCBI database under GenBank accession numbers JYCT00000000, JYCU00000000, and JYCV00000000, respectively.

The genomes of CU_GBS_98 and CU_GBS_08 were completed (GenBank Accession nos.: CP010875 and CP010874 respectively). Draft genome scaffolds were built using the CONTIGuator software (version 2.7.4) [67], with reference to a GBS complete genome (NEM316, accession no. NC_004368). Gaps between adjacent contigs were defined using the Geneious (version R6.1.5, http://www.geneious.com) and Mauve software (using progressive Mauve aligner, version 2.3.1) [68]. All gaps were successfully closed with PCR, and the complete genomes of CU_GBS_98 and CU_GBS_08 were deposited in the NCBI database.

Both draft and complete genomes were mapped to the reference genome of CU_GBS_08 to call small indels and SNPs using the nucmer script (--mum and --breaklen=500) from the MUMmer software (version 3.23) [69]. Functional effects of the identified indels (in-frame or frame-shift indels) and SNPs (synonymous/non-synonymous/stop-codon mutations) were determined according to gene annotations of the reference genome.

### Generation of *ΔBceR* mutant strain using allelic replacement

PCR products containing (a) ∼900 bp of sequence upstream of the *BceR* gene, and (b) the last 58 bp of the *BceR* gene to approximately 900 bp downstream of the gene, were amplified by PCR. The fragments were digested by restriction enzyme *EcoRI* and ligated with T4 DNA ligase according to the manufacturer’s protocol (NEB, MA, U.S.A.). The ligated products were amplified by crossover PCR. The PCR product and the thermosensitive plasmid pJRS233 were digested with restriction enzymes *KpnI* and *BamHI,* ligated, and then transformed into XL-Blue competent cells (Agilent, CA, U.S.A.). The resulting plasmid was extracted with the Plasmid Maxi Kit (Qiagen, Limburg, Netherlands) and transformed by electroporation into CU_GBS_08 [70]. Transformants were selected at 30°C with 1 μg/ml erythromycin in Todd Hewitt agar with 0.5% yeast extract and 5% defibrinated horse blood. Cells with the plasmid integrated into the chromosome were selected at 37°C under erythromycin pressure, and subsequently passaged at the same temperature in the absence of erythromycin for plasmid excision.

### Construction of complementation plasmid to rescue *ΔBceR* phenotypes

A plasmid was constructed to express the full-length *BceR* gene, and a 500 bp fragment of the upstream region of *BceR* gene was amplified with primers containing *BamHI* and *Xbal* sites, and was cloned into the *BamHI* and *Xbal* sites of pDL289, to create the expression vector of BceR, pDL289-BceR. Inserts and reading frames were confirmed by sequencing. pDL289-BceR was introduced into *ΔBceR* mutant strain by electroporation.

### MIC determination

The minimum inhibitory concentration (MIC) of antimicrobial agents was determined by the microbroth dilution method, according to the Clinical and Laboratory Standards Institute (CLSI, 2011).

### RNA extraction and real time-PCR

*S. agalactiae* was plated on blood agar plates and incubated at 35°C in 5% CO_2_. Sub-inhibitory bacitracin (Bac) concentration values were determined by monitoring cell growth in THB with or without a range of bacitracin in 96-well plates. In brief, overnight culture cells were resuspended and adjusted to an OD_600_ 0.8. One percent bacterial suspension was prepared to obtain a final inoculum of 1×10^6^ to 5×10^6^ CFU per well in 200 μl THB with or without Bac at 1/2, 1/4, and 1/8×MICs. The bacterial cells were then incubated at 37 °C, and OD_595_ was measured every 30 min using a DTX 880 microplate reader (Molecular Devices, CA, U.S.A.) over 24 h. The minimum bacteria concentration that did not alter the bacterial growth curve was determined as the sub-inhibitory concentration for the described experiment.

Total RNA was extracted with Trizol [71]. Briefly, 2 ml mid-log phase cells were harvested by centrifugation at 6000×g at 4 °C for 10 min. The pellets were resuspended in TE buffer and incubated with 400 μl lysozyme (prepared in TE buffer) (Sigma, MO, U.S.A.) at 37 °C for 30 min. The lysate was treated with 30 μl of 3 M sodium acetate (Sigma, MO, U.S.A.), 90 μl 10% SDS (Merck, Gernsheim, Germany), and 1 ml Trizol (Life Technologies, CA, U.S.A). This was followed by a 5 min incubation at RT before adding 200 μl of chloroform (Merck, Gernsheim, Germany) for 2 min. All of the samples were centrifuged at 12,000xg at 4 °C for 15 min. The supernatant was transferred to a new tube with 1 ml isopropanol (Merck, Gernsheim, Germany) for RNA precipitation. After 2 h of incubation at −20 °C, the tubes were centrifuged at 12,000 g at 4 °C for 15 min with the supernatant discarded. An equal volume of cold absolute ethanol (Merck, Gernsheim Germany) was added to the tube, and centrifuged at 12,000xg at 4 °C for 5 min to obtain the RNA pellet. The pellet was resuspended in 100 μl of DNase-free and RNase-free water. and additionally treated with 2 U of DNase I (Promega, WI, U.S.A.) followed by a 37 °C incubation for 20 min. The RNA quality and quantity were quantified by Nanodrop 1000 (Life Technologies, CA, U.S.A.), and then stored in 20 μl aliquots at −80 °C.

Two hundred nanograms of total RNA of each sample were subjected to cDNA synthesis using a TURBO DNA-free Kit (Applied Biosystems, MA U.S.A.) and SuperScript III Reverse Transcriptase (Invitrogen, CA, U.S.A.) according to the manufacturer’s protocol. Real-time PCR was performed using SYBR Green PCR Master Mix (Invitrogen, CA, U.S.A.) according to the manufacturer’s protocol, and carried out in an ABi 7500 Real-Time PCR Detection System (Applied Biosystems, MA, U.S.A.). Each sample was run in triplicate with 300 nM of each primer (Table S2) under the following conditions: 95 °C for 10 min, 40 cycles of 95 °C for 30 s, and then 60 °C for 1 min. Melting curves were generated by a cycle of 95 °C for 1 min and 60 °C for 1 min. The relative quantitation of mRNA expression was normalized to the constitutive expression of the 16S rRNA housekeeping gene and calculated by the comparative ΔΔCT method.

### H_2_O_2_ stress assay

*S. agalactiae* strains were plated on blood agar plates and incubated at 35 °C in 5% CO_2_. Bacterial cells were suspended with pre-warmed THB with shaking at 200 rpm overnight. The overnight bacterial cells were then 1:100 diluted in THB and incubated at 37°C with shaking at 200 rpm to OD_600_=0.8-1.0. The bacteria were resuspended in THB at a concentration of 4*10^^7^ cfu/ml, and then 40 mM H_2_O_2_ was added at RT for 15 min. After treatment, fresh THY broth was added to stop the reaction, and the bacteria were harvested by centrifugation at 4000xg for 15 min. Bacterial viability after H_2_O_2_ treatment was then examined by culture and enumeration of the bacterial colonies. Serial dilutions of the medium were conducted for cfu counting. Each experiment was conducted in triplicate.

### Determination of biofilm biomass by crystal violet staining and cfu counting

*S. agalactiae* strains were plated on blood agar plates and incubated at 35 °C in 5% CO_2_. Overnight bacterial cells were suspended with pre-warmed THB overnight. 24-well flat bottom plates (Costar, MA, U.S.A.) were used to support the biofilm growth. Then, the overnight bacterial cells were 1:100 diluted in THB and incubated at 37°C with shaking at 200 rpm to OD_600_ 0.8-1.0. The bacteria were harvested by centrifugation at 4000x g for 15 min. After washing with PBS, the cells were diluted 1:10 with pre-warmed THB, and 500 μl cells were added to each well of 24-well plate and incubated at 37°C, 5% CO_2_ overnight without shaking. All samples were run in triplicate.

Biofilm biomass was quantified by measuring the absorbance of crystal violet [55]. After removing the culture medium, the plates were gently washed with PBS twice to remove the floating cells. Biofilms were stained with 300 μl of 0.5% crystal violet (Sigma, MO, U.S.A.) (prepared in 10% ethanol) for 15 min at RT. After staining, the plates were gently washed with PBS three times and dried at RT. 500 μl 95% ethanol was then added to each well and incubated for 15 min to dissolve the biofilms. The OD_595_ values were measured using a DTX 880 plate reader (Molecular Devices, CA, U.S.A.).

Bacterial viability in biofilms was also examined by enumeration of bacterial colonies. After removing the culture medium, the plates were gently washed with PBS twice to remove the floating cells, followed by addition of 500 μl fresh THB to each well. The cells were collected by scraping the bottom of each well with a sterile cell scraper. Serial dilutions of the medium were conducted for cfu counting. Each experiment was conducted in triplicate.

### Determination of biofilm formation by confocal laser scanning microscope

Biofilms were grown on 8-well chamber slides (Nunc, IL, U.S.A.) for visualization under confocal laser scanning microscope. Biofilms were gently washed with PBS three times and stained with the fluorescent dye LIVE/DEAD BacLight bacterial viability kit 7002 (Life Technologies, CA, U.S.A.) for 15 min at RT. The dye was discarded, and the cells were washed twice with PBS and fixed with 4% neutral buffered formalin for 30 min at RT. After removing the fixative and washing three times, cells were removed from the wells to the slide, followed by the addition of 100 μl saline to each well of the chamber to prevent air bubbles when placing a coverslip on the gasket. The edges of the cover slides were sealed and dried for at least 1 h at RT before confocal images were taken and Z-stacks were captured with a Zeiss LSM 880 system (Carl Zeiss, Jena, Germany). Stained slides were protected from exposure to light. The assays were performed at least three times.

### Mitogenicity and cytokines release in human lymphocytes

#### Lymphocyte proliferation assay

Bacteria were grown in Todd-Hewitt broth (Oxoid) with 0.2% yeast extract overnight at 37°C. The overnight cultures were diluted 1:100 in fresh THB, grown to mid-log phase, harvested by centrifugation at 3000xg for 10 min, and then washed three times with phosphate-buffered saline (PBS). Pelleted cells were resuspended in PBS, heat-killed (100°C, 30 min), and underwent centrifugation at 11,000 g for 20 min at 4°C to remove cell debris. Supernatant (GBS cell extract) was aliquoted and stored at −80°C until required. Protein concentrations were determined by protein assay dye reagent concentrate (Biorad) using bovine serum albumin (Sigma) as a standard.

Peripheral blood mononuclear cells (PBMCs) were isolated from whole blood of healthy individuals (obtained from the Hong Kong Red Cross Blood Transfusion Service) by density gradient centrifugation using Ficoll-Paque (GE Healthcare). The human mononuclear cells were washed with PBS, resuspended in medium (RPMI 1640 with 10% FBS), and seeded at 2 × 10^5^ per ml in a 96-well View Plate (Perkin Elmer). Twenty-four hours later, GBS cell extract (prepared in the first paragraph) at a final concentration of 25 μg/ml was added. Phytohemagglutinin (PHA, 10 μg/ml) and culture medium alone were included as controls. After incubation for 24 h, proliferation of lymphocytes was then detected using alamarBlue (Life Technologies) according to the manufacturer’s protocol. Fluorescence emission was measured using an EnSpire Multimode Plate Reader (Perkin Elmer) at 585 nm with an excitation wavelength of 570 nm. Experiments were performed in triplicate.

#### Cytokines measurement

After stimulation of PBMCs, supernatant from the cell cultures at 3 h, 6 h, 12 h, and 24 h incubation was collected for measurement of cytokine release. Interleukins (IL) −1β, IL-6, IL-8, IL-10, IL-12, and tumour necrosis factor alpha (TNF-α) were evaluated by the ELISA method according to the manufacturer’s instructions (BD Biosciences). Measurements were read at OD_450_nm (EnSpire Multimode Plate Readers, PerkinElmer).

#### Murine infection model

Animal experiments were performed with permission of the Animal Experimentation Ethics Committee (AEEC) of the Chinese University of Hong Kong.

The virulence of GBS III-4 mutant strains *ΔBceR* was tested against the wild type strain CU_GBS_08, the CU_GBS_12 strain with natural truncations in *BceR,* and a non-invasive ATCC 12403 control strain in a mice model. The GBS inoculum was prepared by 1:100 dilutions of overnight cultures into Todd Hewitt broth. Cultures were incubated at 37°C, and then bacteria cells were harvested by centrifugation at 2500 rpm for 10 min at 4°C. The pellet was washed twice and resuspended in 5 ml phosphate buffered saline (PBS). GBS inoculums were prepared by serial dilution of the PBS suspension from 10^9^ to 10^5^ CFU/ml. Dilutions were confirmed by colony counts on blood agar. Six-week-old CD1 mice were purchased from The Laboratory Animal Services Centre (The Chinese University of Hong Kong, Hong Kong). The mice were infected by intraperitoneal injection with 0.1 ml GBS inoculums from 10^9^ to 10^5^ CFU/ml. The control group was injected with an equivalent volume of sterile PBS. Each group contained five mice. The survival rate within 10 d post-infection was used to calculate the median lethal dose (LD_50_) [72].

#### 2-D Page and mass spectrometry analysis

*S. agalactiae* strains were plated on blood agar plates and incubated at 35 °C in 5% CO_2_. Bacterial cells were suspended with pre-warmed THB with shaking at 200 rpm overnight. Then the overnight bacterial cells were 1:100 diluted in THB and incubated at 37°C with shaking at 200 rpm to mid-log phase, and then the bacterial cells were harvested by centrifuging at 4000g for 20min at 4°C. For whole protein extraction, instruction of the total protein extraction kit (Bio-Rad, USA) was followed, and protein Quantitation was done using RC DC Protein Assay reagent (Bio-Rad, USA). The protein sample mixture was applied onto the rehydration tray (Bio-Rad, USA), and then a 11cm pH 4-7 IPG strip was placed on top of the sample, with the gel surface facing the sample mixture. The strip was covered with mineral oil (Bio-Rad, USA), and leave at room temp for 12hrs-16hrs without disturbance. The rehydrated strip was rinsed with fresh mineral oil and placed on the well of the focusing tray (Bio-Rad, USA), with gel surface facing the well bottom and the well was covered with mineral oil, then the focusing tray was placed to the isoelectrofocusing machine for isoelectrofocusing (IEF). After isoelectrofocusing, the strip was rinsed with Milli Q water to remove the mineral oil, and equilibrated in freshly prepared equilibration buffer A with 1% DTT (w/v) for 15mins, followed with freshly prepared equilibration buffer B with 4% IAA (w/v) for 15mins. After equilibration, the gel was rinsed with MOPS buffer (Bio-Rad, USA), and then placed on the well of a 10% Bis-Tris XT Criterion gel, and the gel was placed into the electrophoresis tank for electrophoresis. After electrophoresis, the 2-DE gel was stained with Colloidal Blue (Bio-Rad, USA), following the instruction of the kit.

The gel photos were normalized and compared using software PDQuest (Version8.0.1, Bio-Rad, USA), Boolean method was chosen for detecting the proteins with statistic significantly difference in expression between GBS III-4 wild type and mutant strains. From the results we found that 3 proteins have significantly decreased expression in *ΔBceR* mutant strain (>2 folds reduction in expression), and boolean operation was used for comparation of the intensity of the expression of protein spots. These three protein spots were cut and prepared for further identification using mass spectrometry. Finally, the selected protein spots were cut from the original 2-DE gel and to a proteomic core laboratory of The University of Hong Kong for Mass spectrometry analysis.

#### Statistical analysis

Data were expressed as mean ± SD. Statistical comparisons between different treatment groups were carried out using one-way analysis of variance (ANOVA), followed by *post-hoc* Dunnett’s test using GraphPad Prism 6.05 for Windows (GraphPad Software, San Diego CA, U.S.A.). Differences were considered as significant at *p* < 0.05, and were denoted as \**p* < 0.05, \*\**p* < 0.01, \*\*\**p* < 0.001, and \*\*\*\**p* < 0.0001.

## Acknowledgement

We are grateful to those that performed critical readings of the manuscript. We thank Professor Craig E. Rubens for kindly providing the plasmid pJRS233 used for *BceR* knock out. We are also grateful to Professor Marc Ouellette for generously providing the plasmid pDL289 used for *BceR* complementation.

## Supporting Information Legends

**S1 Table.**
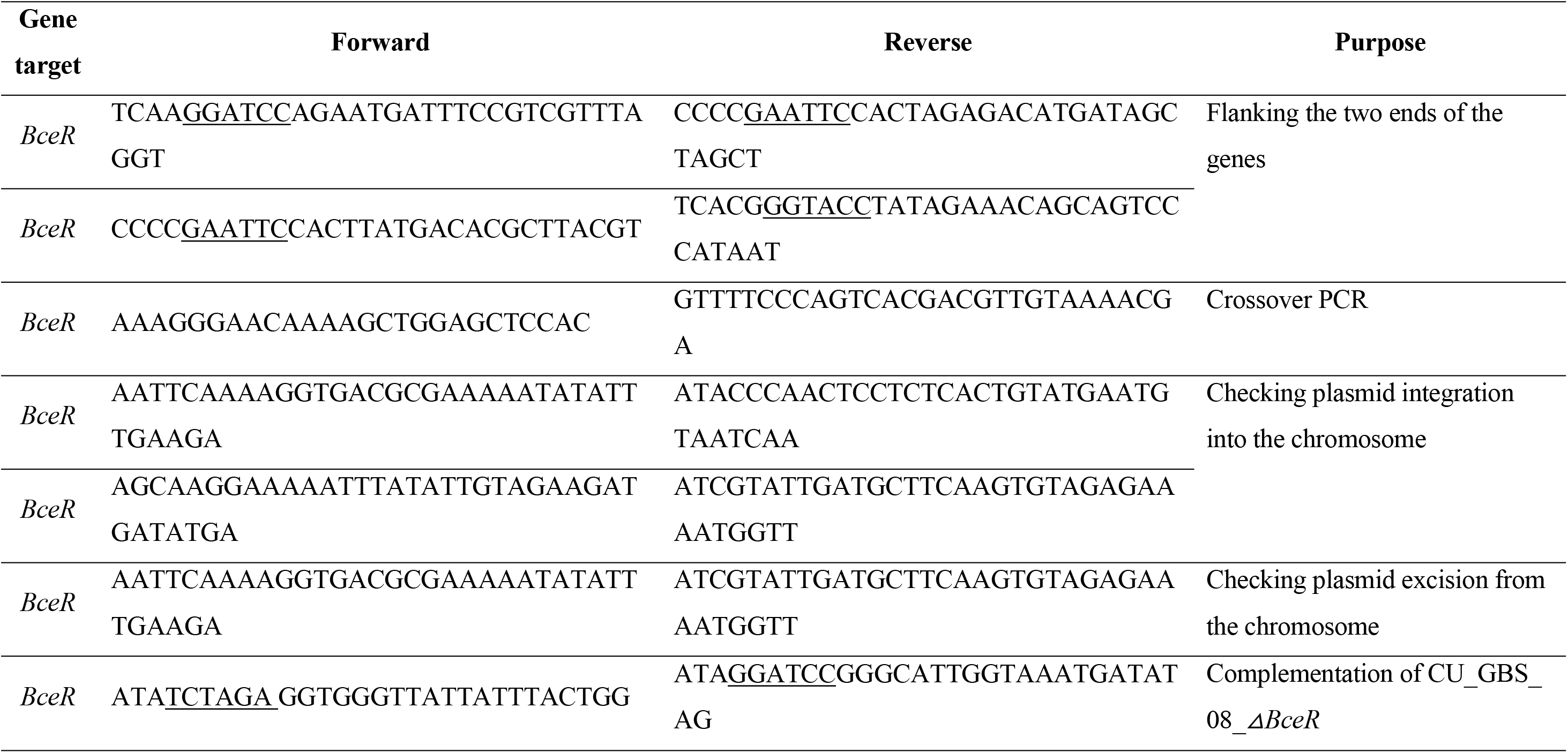
List of primers for constructing *ABceR* mutant and complementary strains.

**S2 Table.**
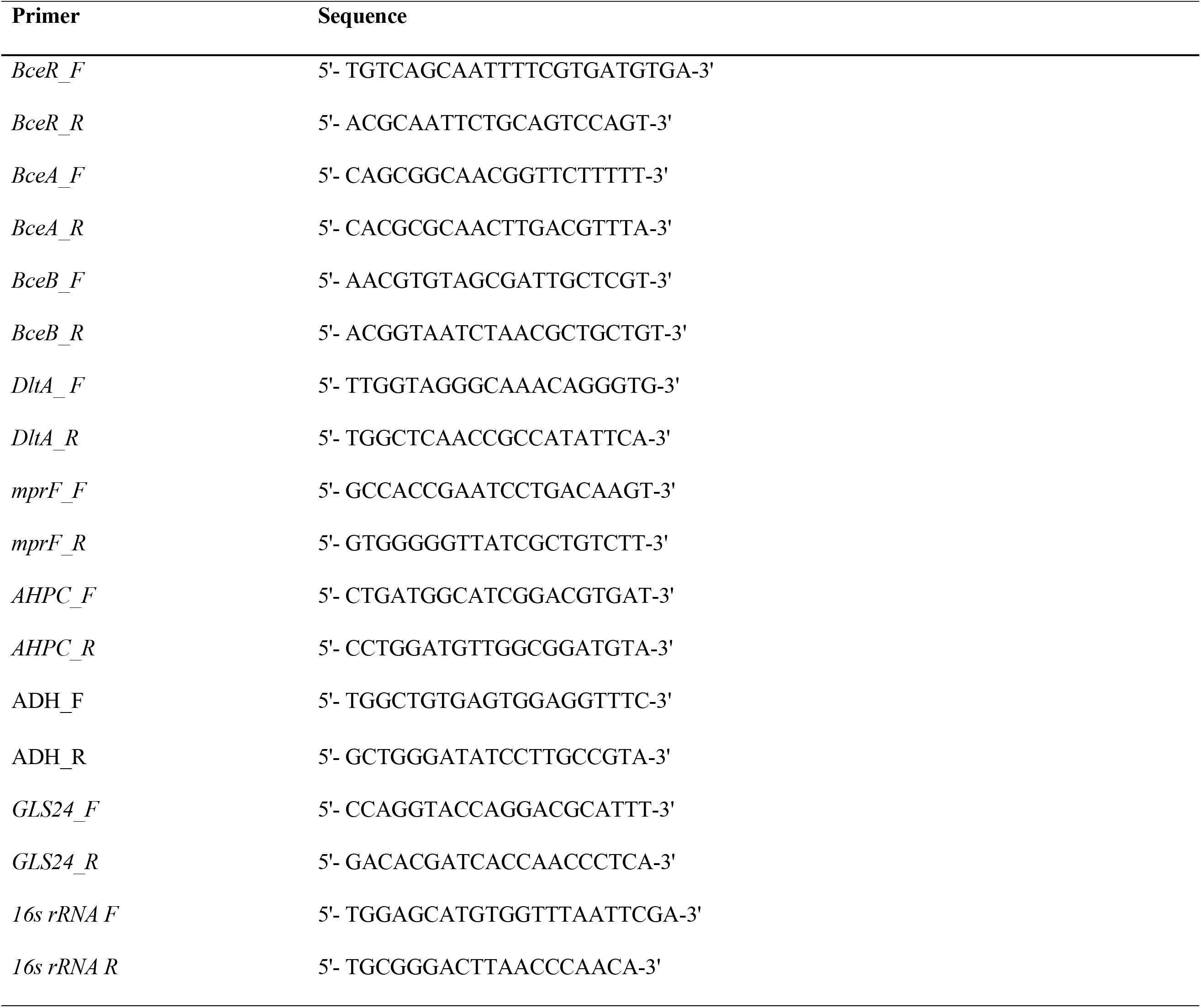
Primers used in real-time PCR.

